# The Garden of Forking Paths: Reinterpreting Haseman-Elston Regression for a Genotype-by-Environment Model

**DOI:** 10.1101/2023.12.19.572468

**Authors:** Guo-Bo Chen

## Abstract

Haseman-Elston regression (HE-reg) has been known as a classic tool for detecting an additive genetic variance component. However, in this study we find that HE-reg can capture GxE under certain conditions, so we derive and reinterpret the analytical solution of HE-reg. In the presence of GxE, it leads to a natural discrepancy of linkage and association results, the latter of which is not able to capture GxE if the environment is unknown. Considering linkage and association as symmetric designs, we investigate how the symmetry can and cannot hold in the absence and presence of GxE, and consequently we propose a pair of statistical tests, Symmetry Test I and Symmetry Test II, both of which can be tested using summary statistics. Test statistics, and their statistical power issues are also investigated for Symmetry Tests I and II. Increasing the number of sib pairs is important to improve statistical power for detecting GxE.

## Introduction

To investigate linkage between a quantitative trait and a marker locus, Haseman and Elston proposed the seminal “Haseman-Elston regression” (HE-reg) method (Haseman and Elston, 1972). Since its inception, HE-reg has been a rich garden, which fertilizes many other methods and leads to various improvements (Elston *et al*., 2000). It has been adapted as a major tool for linkage scans (Cardon and Fulker, 1994; Fulker and Cardon, 1994; Xu *et al*., 2000; Sham and Purcell, 2001), or for estimating heritability at a genome-wide scale using sib pairs (Visscher *et al*., 2006). Since the rise of genome-wide association studies, HE-reg has been reinvigorated for addressing the problem of “missing heritability” (Yang *et al*., 2010). Through the linkage kernel (identical-by-descent, IBD) of HE-reg being replaced by IBS (identity-in-state, IIS, also known as IBS), HE-reg can estimate SNP-heritability with good accuracy (Chen, 2014; Golan *et al*., 2014; Bulik-Sullivan, 2015; Wu and Sankararaman, 2018). Recently, because of the possibility of accumulating new pedigree data (Kong *et al*., 2018; Kaplanis *et al*., 2018), HE-reg has been further modified to deal with linkage analysis at an unprecedented scale (Young *et al*., 2018; Zajac *et al*., 2023).

Due to technical difficulties, there has been some debate on the properties of HE-reg (DeFries, 2010). Nevertheless, there is a consensus that HE-reg is a model-free framework to detect an additive genetic variance component. We here report a long overlooked ambiguity that HE-reg captures if there is any genotype-environment (GxE) variance component besides the additive genetic variance. Following the original approach of HE-reg, we derive the analytical result for the regression coefficient of HE-reg when GxE is present. Due to its historical importance, this reinterpretation for HE-reg will shed light on a forking path in the garden.

This study is organized around two kinds of heritability: *h*^2^, which is consistent with the conventional definition heritability, and 𝒽 ^2^, which additionally includes GxE. We briefly review HE-reg and introduce GxE for a labile trait of that is subject to the norm of reaction. GxE is found to be naturally integrated into HE-reg without the need to include any additional parameters, and this leads to a new interpretation of HE-reg. As HE-reg has been widely used for both linkage and association studies, either in a single-locus or whole-genome context (Sham and Purcell, 2001; Visscher *et al*., 2006; Chen, 2014), we here pursue symmetric and asymmetric forms between linkage and association studies under the new interpretation of HE-reg. For linkage and association, two pairs of symmetries are developed: Symmetry I for single-locus estimation of heritability, and Symmetry II for whole-genome estimation of heritability. These two symmetry groups are represented by their respective test statistics – *t*-tests here. On the basis on the good analytical properties of HE-reg, we envisage linkage and association in a complementary manner to each other and empower the investigation of genetic architecture. When there is GxE, these two symmetries should not hold, but a large sample size—especially for the size for sib pairs—is needed to detect possible asymmetry between linkage and association studies.

## Methods

We adapt the original sib pair design of HE-reg, and have a linear model for the additive effect of a casual variant

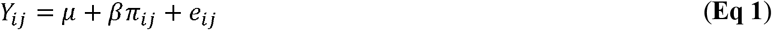

in which *Y*_*ij*_ = (*y*_*i*_ − *y*_*j*_)^2^ for a pair of individuals *i* and *j* who are a sib pair *μ*, the mean of the model, *β* the regression coefficient *π*_*i j*_, the identity-by-descent (IBD) of the sib pair, and *e*_*i j*_ the residual. We consider *K* pairs of sibs from *K* independent families, so *Y*_*ij*_ is a *K* × 1 vector. Now we take a close look at its signature result, the interpretation of the regression coefficient *β*, which is written as

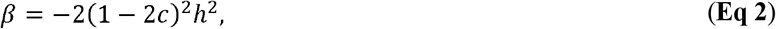

in which *c* is the recombination fraction between the marker and a causal variant, and *h*^2^ the variance explained by the causal variant. **Eq 2** is the commonly accepted standard interpretation for HE-reg (Lynch and Walsh, 1998). Here we have not distinguished *h*^2^ from 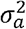 (additive genetic variance of a locus), which can be reconciled by scaling. In the context below, we choose the appropriate notation. *h*^2^ or 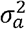.

### Norm of reaction for a labile trait

Sibling differences such as one drinks but the other not makes a form of personalized “environment”, but we only focus on environmental factors that can apply to the whole family, such as highland habitants vs lowland habitants. A typical non-removable GxE is present if both genotype effects “crossover” (Wang *et al*., 2010), as the norm of reaction defined below:

#### Scheme I (highland habitants)

genotypes *BB, Bb* and *bb* with their corresponding effects *G*_*BB*_, = *a,G*_*Bb*_=*d* and *G*_*bb*_ = −*a*;

#### Scheme II (lowland habitants)

genotypes *BB, Bb* and *bb* with their corresponding effects *G*_*BB*_ = −*a, G*_*Bb*_*= d* and *G*_*bb*_ *=a*

We assume *K*=*K*_1_ + *K*_2_ sib pairs, and *K*_1_ sib pairs are highland habitants and *K*_2_ sib pairs lowland. Now, we have. **Table 1**, which is a modification of the original HE paper (please refer to “Table I Conditional Distribution of *Y*_*j*_” in Haseman and Elston’s original paper). According to **Table I**, under **scheme I**, we have

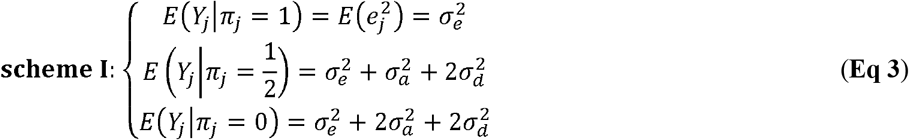

in which 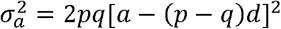 and 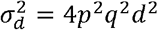.

**Table 1.** A rework of the original Table I for HE-reg when the target locus harbors genotype-environment interaction.

| Sib pair | $Y_{ij} = (y_i - y_j)^2$ | | Conditional probability for IBD ($\pi_j$) | | |
| --- | --- | --- | --- | --- | --- |
| | scheme I ($\omega_1$) | scheme II ($\omega_2$) | $\pi_j = 0$ | $\pi_j = \frac{1}{2}$ | $\pi_j = 1$ |
| | $G_{BB} = a, G_{Bb} = d, G_{bb} = -a$ | $G_{BB} = -a, G_{Bb} = d, G_{bb} = a$ | | | |
| $BB - BB$ | $e_j$ | $e_j$ | $p^4$ | $p^3$ | $p^2$ |
| $bb - bb$ | | | $q^4$ | $q^3$ | $q^2$ |
| $Bb - Bb$ | | | $4p^2q^2$ | $pq$ | $2pq$ |
| $BB - Bb$ | $(a - d + e_j)^2$ | $(a + d + e_j)^2$ | $2p^3q$ | $p^2q$ | 0 |
| $Bb - BB$ | $(-d + a + e_j)^2$ | $(-a - d + e_j)^2$ | $2p^3q$ | $p^2q$ | 0 |
| $Bb - bb$ | $(a + d + e_j)^2$ | $(a - d + e_j)^2$ | $2pq^3$ | $pq^2$ | 0 |
| $bb - Bb$ | $(-a - d + e_j)^2$ | $(-d + a + e_j)^2$ | $2pq^3$ | $pq^2$ | 0 |
| $BB - bb$ | $(2a + e_j)^2$ | $(2a + e_j)^2$ | $p^2q^2$ | 0 | 0 |
| $bb - BB$ | $(-2a + e_j)^2$ | $(-2a + e_j)^2$ | $p^2q^2$ | 0 | 0 |

**Notes:** This table is adapted from Table I from the original HE-reg paper (Haseman and Elston, 1972), but we expand the column “ $Y_{ij} = (y_i - y_j)^2$ ” to have GE shown as **scheme I** and **scheme II**. $\omega_1 = \frac{K_1}{K_1 + K_2}$ the proportion of sib pairs having **scheme I** genetic effects, and $\omega_2 = \frac{K_2}{K_1 + K_2}$ **scheme II**.

Now turn to **scheme II**,

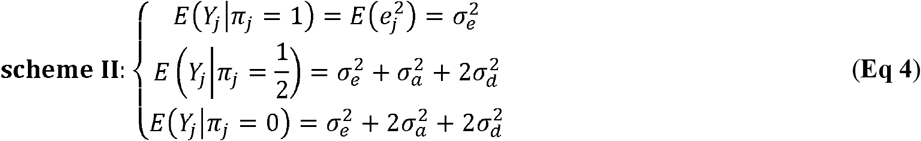

Here we have 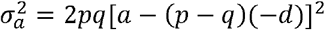 which leads to the identical result like **Eq 3**. As in the original HE paper,we drop off 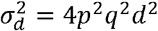 because of ignorable dominance genetic variance, and **Eq 4** is identical to the **Eq 3**, indicating that HE-reg does not distinguish GxE, and consequently does not distinguish GxE from a pure additive model. So, we arrive at a generalized expression for the HE-reg regression coefficient:

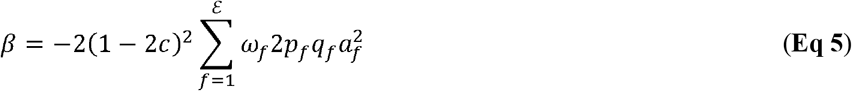

in which *ε* is the number of environments. **Eq 5** actually slices the original HE-reg into much thinner layers, each of which covers a fraction *ω*_*f*_ of sib pairs, who are sampled from environment *f*. We assume *a*_*k*_ follows *a*_*k*_, ∼ *N* (*a,E*^2^), GxE in which *a* the inter-family additive effect (marginal effect) and *E*^2^is the variation due to familial origin – an analogue of 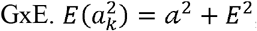, and **Eq 5** can be written in an even more general form as

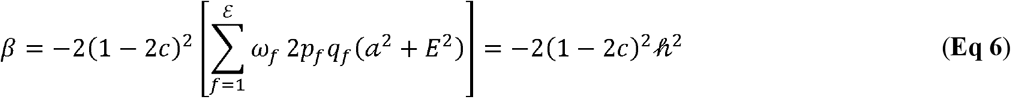

Even though the model is identical to the original HE-reg, **Eq 6** indicates that 𝒽^2^ has an additive variance component, which can be detected via an additive effect model, together with a GxE component. As discovered by recent large-sample GWAS, causal variants are very saturated (Yengo, 2022). We now omit the factor (1 −2*c*)^2^ in **Eq 6** in the text below. The complexity of the original HE-reg paper arises largely to deal with(1 −2*c*)^2^.

In particularly, for HE-reg we have its null and alternative hypotheses:*H*_0_: *β* = 0;*H*_1_ *β* ≠ 0, so for **Eq 6** we construct a *t*-test *t*_*LS*_ for single-locus linkage analysis:

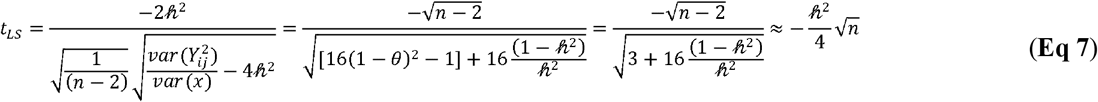

For sib pairs, in *θ* = 0.5 in **Eq 7**, and the approximation holds when 𝒽 ^2^ is small. The detailed elements for constructing **Eq 7** are given in **Table 2**.

**Table 2.** The test statistics for various Haseman-Elston regressions.

| Model | $var(y)$ | $var(x)$ | $\mathcal{B}$ | $\sigma_B$ | $t$-test statistic |
| --- | --- | --- | --- | --- | --- |
| Symmetry I: single-locus model |  |  |  |  |  |
| HE-reg<br>(sib pairs) | $var[Y_{ij}^2] = 8(1 - \theta)^2 h^4 + 8(1 - h^2)$ | $\frac{1}{8}$ | $-2\tilde{h}^2$ | $\sqrt{\frac{1}{(n-2)}} \sqrt{8var(Y_{ij}^2) - 4\tilde{h}^4}$ | $t_{LS} = \frac{-\sqrt{n-2}}{\sqrt{[16(1-\theta)^2 - 1] + 16\frac{(1-h^2)}{h^4}}} \approx -\frac{\tilde{h}^2}{4}\sqrt{n}$ |
| HE-reg<br>(unrelated) | $\sigma_y^2 = 1$ | 1 | $-2h^2$ | $\sqrt{\frac{1}{[\frac{n(n-1)}{2} - 2]}} \sqrt{8 - 4h^4}$ | $t_{AS} = -\frac{\sqrt{\frac{n(n-1)}{2} - 2}}{\sqrt{\frac{2}{h^4} - 1}} \approx -\frac{n}{2}h^2$ |
| Symmetry II: whole-genome model (collapsed genetic architecture) |  |  |  |  |  |
| HE-reg<br>(sib pairs) | $var[Y_{ij}^2] = 8(1 - \theta)^2 h^4 + 8(1 - h^2)$ | $\frac{1}{16\mathcal{L}}$ | $-2\tilde{h}^2 = -2 \sum_k^Q \tilde{h}_k^2$ | $\sqrt{\frac{1}{(n-2)}} \sqrt{16\mathcal{L}var(Y_{ij}^2) - 4\tilde{h}^4}$ | $t_{LW} = \frac{-\sqrt{n-2}}{\sqrt{16\mathcal{L} \left[ 2(1-\theta)^2 + 2\frac{(1-\tilde{h}^2)}{\tilde{h}^4} \right] - 1}}$ |
| HE-reg<br>(unrelated) | $var[Y_{ij}^2] = 8$ | $\bar{\rho}_{mm}^2$ | $-2\tilde{h}^2 = -2 \sum_k^Q h_k^2$ | $\sqrt{\frac{1}{[\frac{n(n-1)}{2} - 2]}} \sqrt{\frac{8 - 4\tilde{h}^4 \bar{\rho}_{mm}^2}{\bar{\rho}_{mm}^2}}$ | $t_{AW} = -\frac{\sqrt{\frac{n(n-1)}{2} - 2}}{\sqrt{\frac{2}{\tilde{h}^4 \bar{\rho}_{mm}^2} - 1}} \approx -\frac{n}{2}\tilde{h}^2\sqrt{\bar{\rho}_{mm}^2}$ |

Notes:
$t = \frac{\mathcal{B}}{\sigma_B} = \frac{\mathcal{B}}{\sqrt{n-2} \sqrt{\frac{var(y)}{var(x)} - \mathcal{B}^2}}$ ; $var(Y_{ij}^2) = 8(1 - \theta)^2 h^4 + 8(1 - h^2)$ , see **Appendix II**; $E(\theta) = 0.5$ here for sib pairs. We consider the marker itself is a QTL,
so $\beta = -2[2p_f q_f (a^2 + \epsilon^2)] \cdot \bar{\rho}_{mm}^2 = \frac{1}{m^2} \sum_{l_1}^m \sum_{l_2}^m \rho_{l_1 l_2}^2$ . The derivation of $var(x)$ for HE-reg sib pairs, see **Appendix III**.

Nevertheless, for a single-locus GWAS linear model **y** = *a+b*_*j*_**z**_*j*_*+e*, in which ***z*** is the standardized genotype for the *j*^*th*^ locus, the *t*-test is 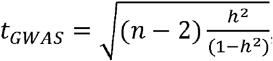, in which *h*^2^ = 2*pqa*^2^ does not include a GxE component. This casts a discrepancy between linkage and association studies.

Replacing IBD with IBS, for unrelated pairs HE-reg is (*y*_*i*_ −*y*_*j*_)^2^= *a* + *𝒷 s*_*ij*_+*e*_*ij*_ .*y*_*i*_ and *y*_*j*_ are the standardized phenotypes for unrelated individuals *i* and *j, s*_*ij*_ the genetic relatedness score, and *e*_*ij*_ the residual. 𝒷 is the regression coefficient. Given an unrelated GWAS sample, the analytical solution for SNP heritability is

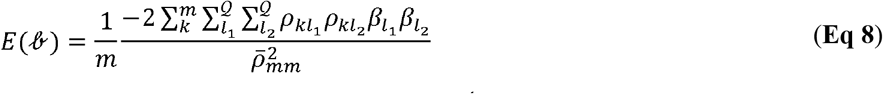

in which 𝒬 is the number of causal variants, often unknown, and 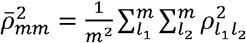. The sampling variance of 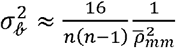 for HE-reg (Chen, 2014).

There are various unexplored paths for possible variation in **Eq 8**, but we here direct our attention to a collapsed form

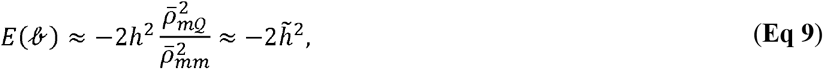

in which 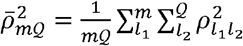 and 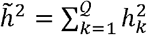 because the summation of the covariances of all pair of variables is zeroed out. A pair of assumptions are needed for **Eq 9**: **I)** the first approximation occurs under the general polygenic assumption that causal variants are randomly allocated along the genome, and **II**) the second approximation further holds when the markers are saturated enough, indicating that nearly every SNP is possibly causal (Yengo, 2022). So, the approximation should be retained for **Eq 8** in view of these two assumptions. Of note, albeit pathologically, counter-examples can be found in which these two assumptions do not hold: see discussion in other studies (Chen, 2014). The corresponding *t*-test *t*_*AW*_ for whole-genome association analysis is (**Table 2**)

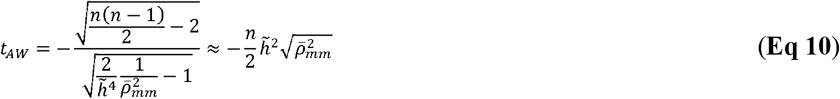

However, when there is only one marker 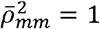, and **Eq 10** leads to a single-locus test for association *t*-test, *t*_*AS*_, which is about (**Table 2**)

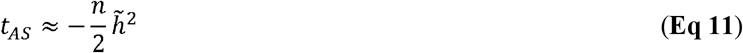

If we replace the π in **Eq 1** with aggregated genome-wide IBD 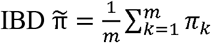, it leads to whole-genome estimation of heritability for a linkage design (Visscher *et al*., 2006):

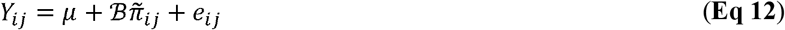

Using the same two assumptions for **Eq 8**, the expectation of 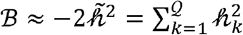, and its sampling variance is about 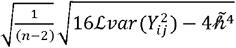, in which ℒ ≈ 32 Morgan is the genetic length of the 22 human autosomes. However, the number of markers in the linkage study determines the sampling variance of ℬ. Consequently, the *t*-test *t*_*LW*_ for whole-genome linkage analysis is (**Table 2**)

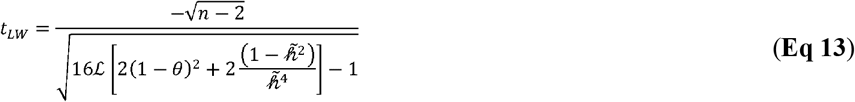

### Pooling all test statistics together

As derived above and summarized in **Table 2**, each *t*-test statistic can be approximately converted to 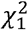 with non-centrality parameter (NCP) of the squared t-test statistic as shown below

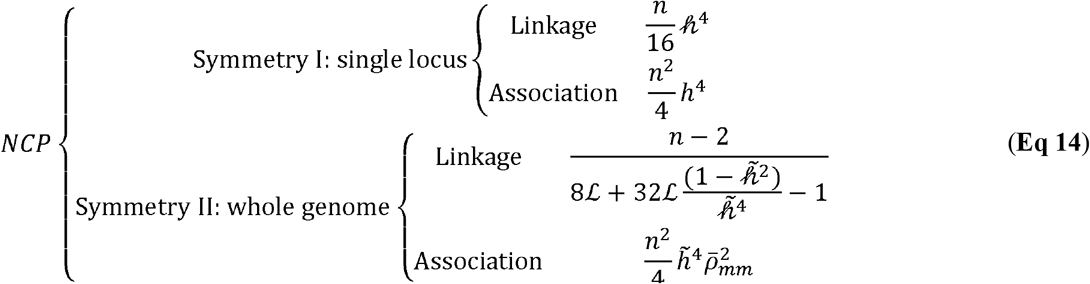

For each paired test statistic, Symmetry I or Symmetry II, under the pure additive model, the test derived from linkage often has less power than that from association; otherwise GxE is available and captured by HE-reg.

### Simulation Results

#### Validation for HE-reg in the presence of GxE

We want to validate the accuracy of the *t*-test statistics in **Eq 7** 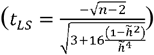in the presence of GxE. We simulated *K* nuclear families, each of which consisted of a pair of unrelated parents and a sib pair. The genotypic effects were set as 1, 0, and -1 for, *BB, Bb*,and *bb* (**Scheme I**) for the first 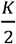 families, and as -1, 0, and 1 for *bb, Bb*, and *BB* (**Scheme II**) for the second 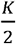 families. The reference allele frequency was uniformly sampled from 0.01 ∼ 0.5; *h*^2^ =0.1, 0.25, and 0.5, respectively; *K =* 200 and 500 respectively. For each simulated sib pair, π (1, 0.5, and 0, and 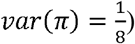 was directly known for the simulation data, and the marker itself was causal so that (1 − 2*c*)^2^ = 1. We resampled each scenario 200 times. As a contrast, for *K* families, we also pooled their 2*K* unrelated parents together, which made a sample for GWAS.

As illustrated in **Figure 1**, since **Scheme I** and **Scheme II** were similar scenarios—but opposite effect size—they had very consistent results within HE-reg. When *K* increased from 200 to 500, the consistency remained. As expected, HE-reg was not as powerful as GWAS in catching a single **Scheme-I or Scheme-II** locus. However, when the **Scheme I** and **II** samples were pooled together and an unremovable GxE was present, HE-reg was more powerful to detect GxE, not just because of the doubled sample size; in contrast, GWAS had no power at all to detect the GxE effect because the effects cancel out under a GWAS model.

**Figure 1.**
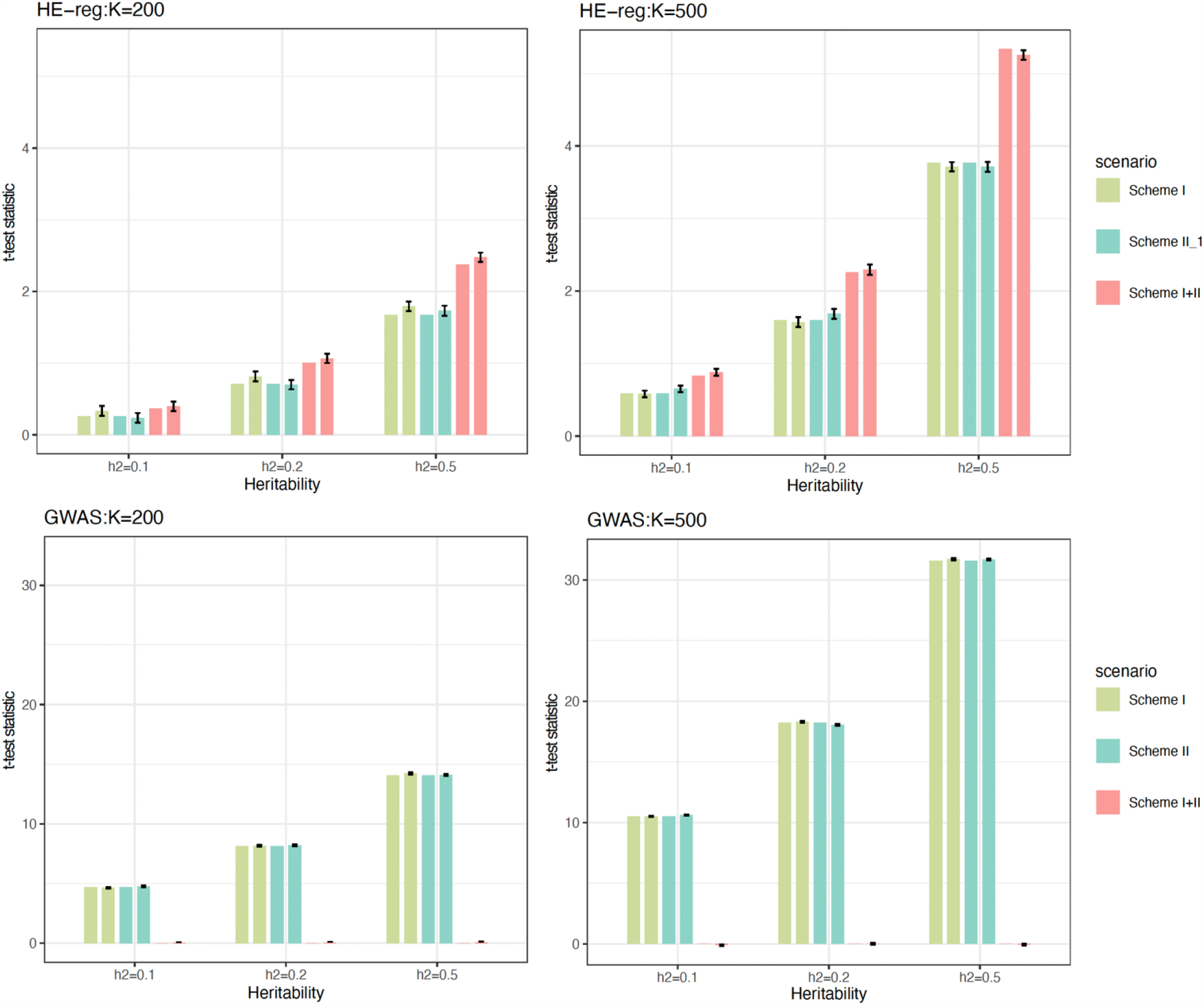
Validation for test statistics for HE-reg and GWAS. 200 simulations for each scenario, and sib pairs were analysed by HE-reg (first row) and their parents were pooled together and analysed by GWAS (second row), such as. In each cluster, the three coloured bars were for scheme I, scheme II, and scheme I+II samples. Of each pair of the same colour bars, the first one was the expected *t*-test statistic calculated by **Eq 7** () for HE-reg (first row) and for GWAS (second row), and the second bar was the mean of the test statistic after 200 simulations and its standard error of mean was represented by the black lines atop. We took absolute values of **Eq 7** and **Eq 8**, and ignored the positive or negative signs.

One criticism might be: why not include GxE in a GWAS model? A GWAS model could be more powerful for detecting a GxE locus, if we knew what the environment was. However, the number of environments is infinite in practice.

### Power calculation for GxE using single-locus HE-reg for a linkage design

It should be noticed that HE-reg has much lower power in linkage than in the comparable association analysis, so it is trivial to compare the statistical power for linkage and association. We considered the statistical power for HE-reg to detect a single locus, which could be in the context of either *h*^2^ or 𝒽^2^. In the simulations, we set the Type I Error rate 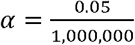 after 1 million genotyped SNPs, and we set 𝒽^2^ = 0.005, 0.0075, 0.01, 0.015 and 0.02, respectively; the total number of sib pairs ranged from 500,000 to 10,000,000, with an increment of 500,000. Given type I error rate 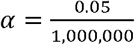 and if the statistical power 0.85 is considered acceptable in practice, for 𝒽^2^ = 0.015, *n* ≈ 3,000,000 sib pairs are required, and for 𝒽^2^ = 0.01, n ≈ 7,000,000 sib pairs are required (**Figure 2**)

**Figure 2.**
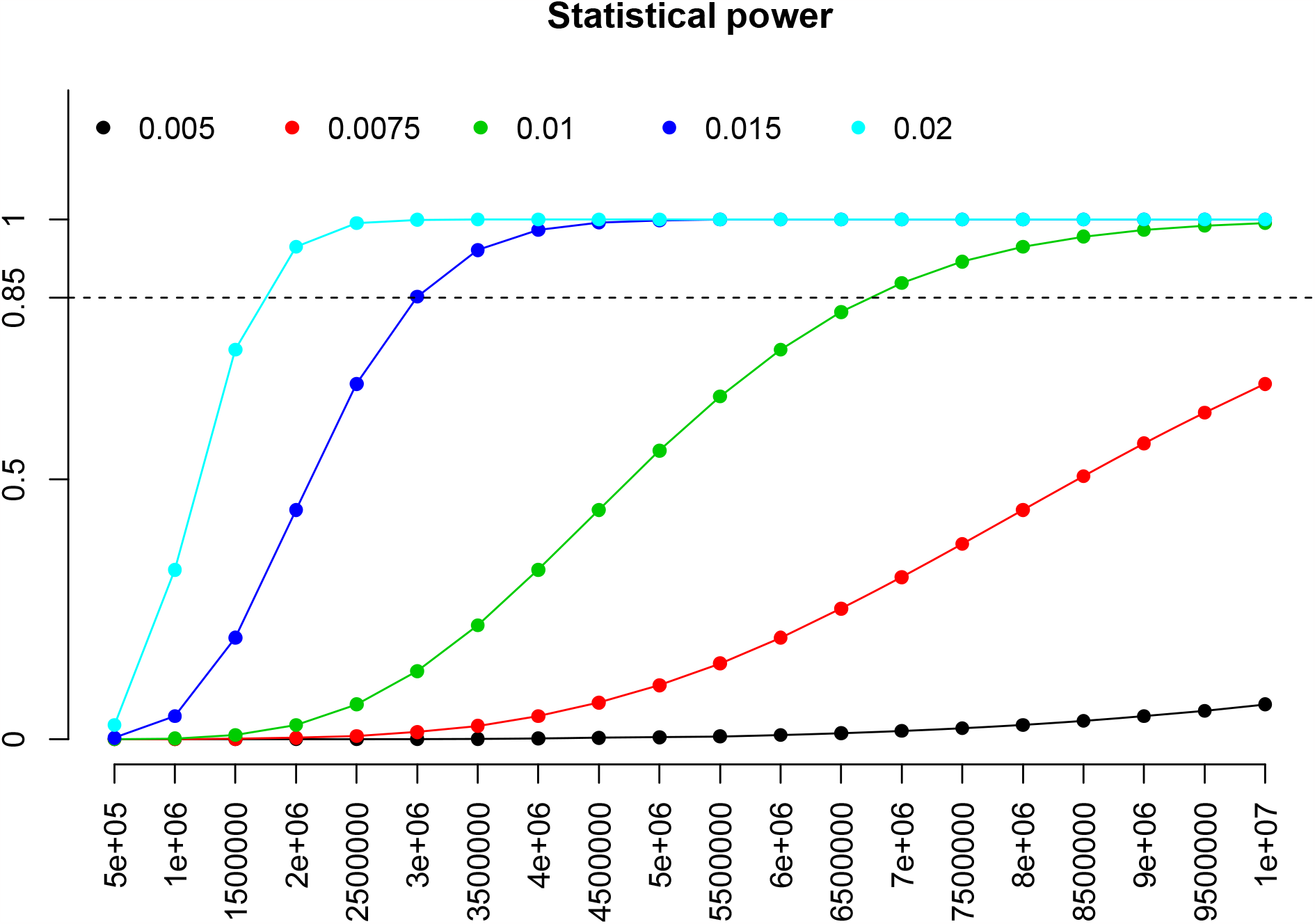
Statistical power to detect a single locus using HE-reg. y-axis represents the statistical power, and x-axis represents the number of sib pairs for HE-reg linkage. Different colour points represent different heritability (𝒽^2^ = 0.005 in black, 𝒽^2^ = 0.0075 in red, 𝒽^2^ =0.01 in green, 𝒽^2^ =0.015 in blue, and 𝒽^2^ = 0.02 in cyan) in cyan). The type I error rate 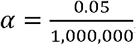, given the number of genotyped SNPs is 1,000,000.

### Statistical Test for Symmetry I

Now we considered **Symmetry I** for a pair of *t*-tests *t*_*LS*_ against *t*_*AS*_ for single-locus tests in linkage and association designs, respectively. The elements for *t*_*LS*_ and 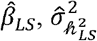and sample size *n*_*LS*_ sib pairs; and for *t*_*AS*_ they are 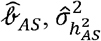, and sample size *n*_*AS*_ (**Table 2**). Using Welch’s m od ifie d two-sample *t*-test we have

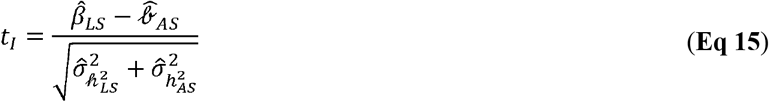

After further approximation 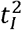 follows 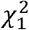 with NCP 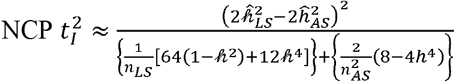. We evaluated statistical power for **Eq 15** by setting a locus with GxE (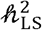 from 0.005 to 0.05) but no additive effect,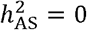, as illustrated in **Figure 3**. The number of SNPs was 1,000,000, and the Type I error rate was 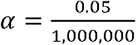. The sib pairs for *n*_*LS*_ =50,000, 150,000, and 300,000, respectively, and *n*_*AS*_ =10,000 and 100,000, respectively. It was obvious that the statistical power benefited from a larger *n*_*LS*_ (**Figure 3**). Of note, Symmetry I could be implemented purely on a pair of summary statistics from single-locus linkage and single-locus GWAS.

**Figure 3.**
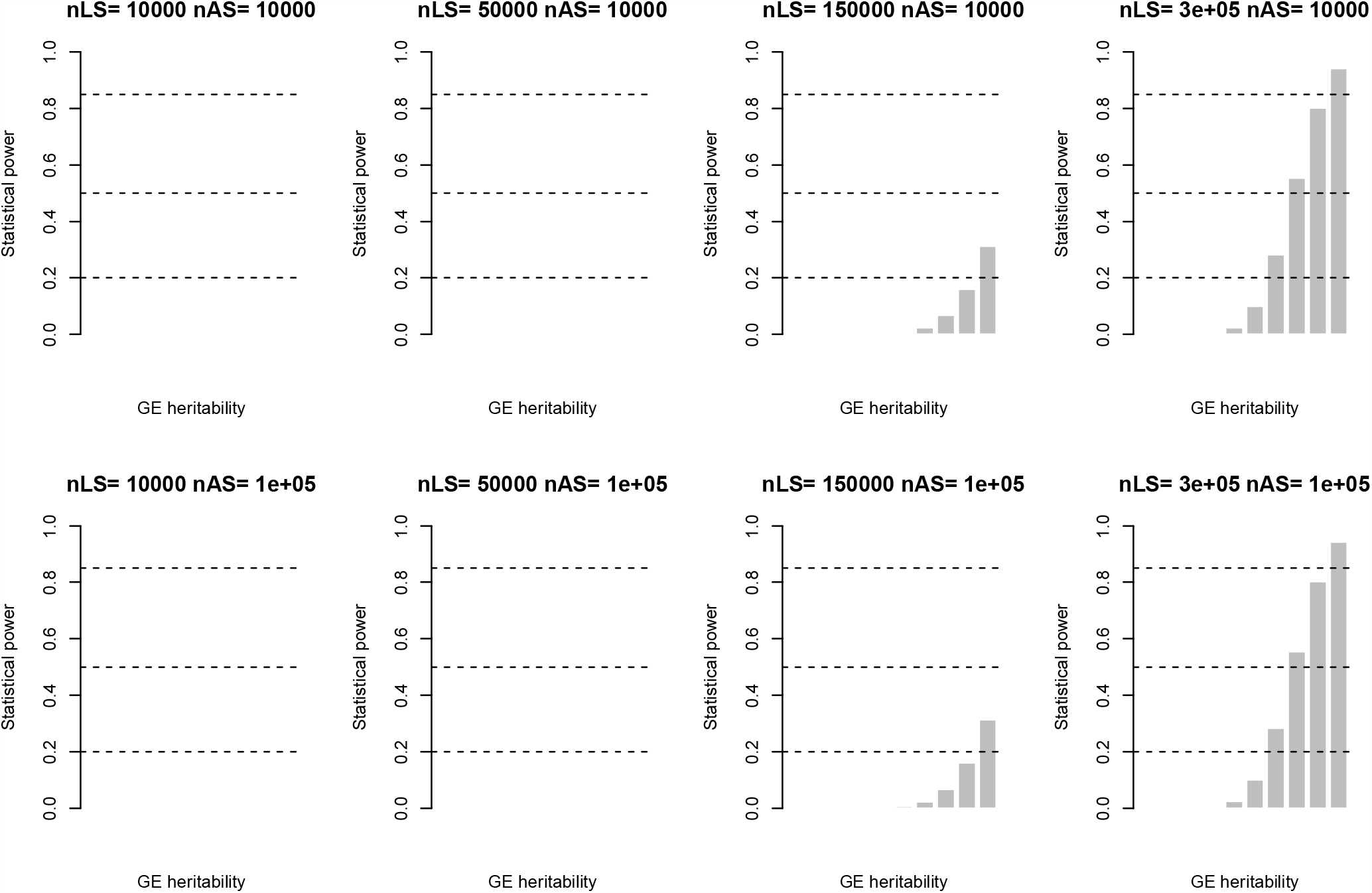
Statistical power for Symmetry I single-locus test for GE. The NCP of the test statistic is 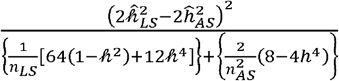 for the transfomred pair of t-tests in **Eq 15**. The y asix is statstical power given 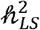 ranged from 0.005 to 0.05 (x-axis), evenly broken into ten values, and 𝒽^2^ =0 in association. The two horizontal lines were for statistical power of 0.25, 0.5, and 0.85, respectively. The sample sizes for linkage (*n*_*LS*_) and for association (*n*_*AS*_) are as shown each subtitle.

**Figure 4.**
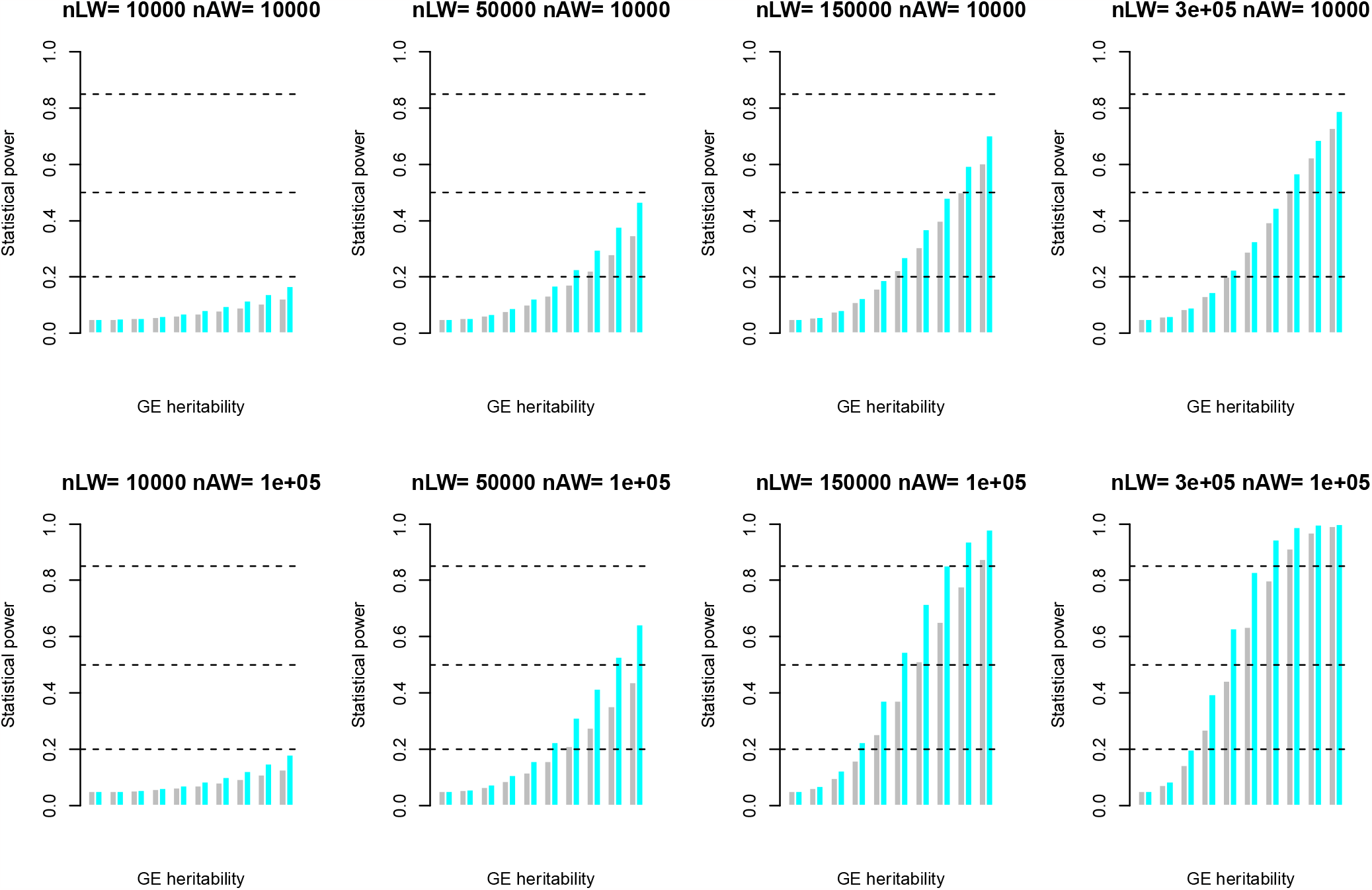
Statistical power for Symmetry II whole-genome test for GE. Numerical evaluation for statistical power of **Eq 16**, and its converted NCP is 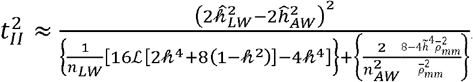. ℒ = 32Morgan, and 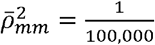. The grey bar is for 𝒽^2^ = 0.25 and 𝒽^2^ = 0.25 + GE, and the cyan bar is for 𝒽^2^ =0.6 and 𝒽^2^=0.6 + GE.CE is always taken value from 0 to 0.2 with increment of 0.02 for each step. The horizontal reference lines are of 0.2, 0.5 and 0.85, respectively in each plot. The sample sizes for linkage (*n*_*LW*_) and for association (*n*_*AW*_) are as shown each subplot.

### Statistical Test for Symmetry II

Analogously, we could test for **Symmet ry II**, a n d it s *t*-test *t* _*II*_ is

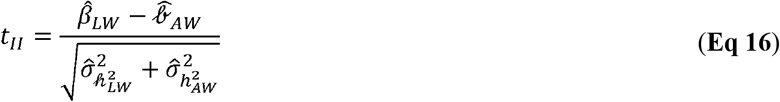

which has NCP 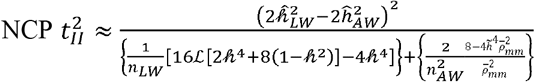. We conducted simulation for **Symmetry II**, and key parameters were set as ℒ = 32 Morgan 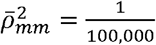 and *h*^2^= 0.25. Two traits were considered, the first trait had and the second *h*^2^ =0.45, which could be detected by whole-genome association HE-reg; and in contrast, with the inclusion of GxE (from 0 to 0.2, with increments of 0.02), its 𝒽^2^ = *h*^2^ + *GxE* which could be detected by whole-genome linkage HE-reg. The sample size for association was *n*_*AW*_= 10,000, and 100,000, respectively, and for linkage it was *n*_*LW*_, =50, 000, 150,000, and 300,000, respectively. The sample size should be larger than 50,000 for linkage, otherwise hardly a sensible difference could be statistically detected.

## Discussion

As revealed in this study, the original HE-reg actually captures not only an additive effect but also GxE. However, a GWAS model is very likely to miss such a GxE location if the environmental variable is unknown. So, given the intriguing behaviour as investigated between HE-reg and GWAS, it naturally leads to the discrepancy for linkage and association studies as long as genes harbor familial variation. The real advantage of our reinterpretation for HE is that a gene desert in GWAS may lead to rich results in single-locus HE-reg linkage, which can detect such a locus without specifying the environment (**Figure 1**). If there are unexplored hotspots for GxE across the genome, it is likely to observe a GxE landscape for various traits and enrich our understanding of genetic architecture. According to our power calculation (**Figure 2**) more sib pairs may enhance the statistical power of HE-reg linkage for detecting GxE. **Symmetry I** and **II** tests can be implemented on summary statistics (**Figure 3** and **4**), so if we have a large sibling data resource, it is possible to construct summary statistic tests for GxE. So far, the publicly available sib pairs may be drawn from UK Biobank (Bycroft *et al*., 2018), but unfortunately there are far less, say about 20,000 sib pairs, than a sample size that leads to practical statistical power.

One of the agenda items in addressing the “missing heritability” question is to narrow down the heritability gap between the family-based studies (linkage) and unrelated GWAS samples (association). In the absence of GxE, increasing the number of variants gives a possible way to solve this issue. The trait most studied is height, the heritability of which is estimated in European descendants using various markers. The benchmark source of family-based heritability estimate of 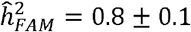 was based on 950 quasi-independent full-sib pairs, and their whole-genome IBD was estimated using 791 autosome microsatellite markers (Visscher *et al*., 2006); the compared SNP-heritability was 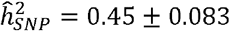, which was estimated from 3,925 unrelated samples and 294,831 SNP markers (Yang *et al*., 2010). The 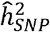 so far has approached 0.68 ± 0.10 using 25,465 unrelated European descendants on the whole-genome sequenced TopMed data with 33.7 × 10^6^variants (Wainschtein *et al*., 2022). Given such a small heritability was sample size for the compared family study, it was not likely to reach statistical significance. **Symmetry II** test indicates another way for interpreting the heritability gap. The gap is not necessarily about inclusion of more variants but rather about using a family-based design instead.

## Acknowledgements

In studying statistical genetics, the author have been helped by Dr Elston much, and this study is consequently dedicated to the semicentennial publication of the original Haseman-Elston regression. HE-reg was based on early works by Lionel Penrose, such as “Genetic linkage in graded human characters” (*Annals of Eugenics*, 1938, 8:233-237) and “The detection of autosomal linkage in data which consist of pairs of brothers and sisters of unspecified parentage” (*Annals of Eugenics*, 1935, 6:133-138). However, in the seminal HE-reg paper, the citation for Penrose’s work is “Penrose, L. S. (1938), Genetic linkage in graded human characters, *Ann. Eugen*. 6:133-138”, an obviously mix-up of these two papers. “The garden of forking paths” is a short story written by Jorge Luis Borges.

## Appendix I Least squares estimation

To a linear model of *n* observations

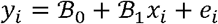

Its least squared estimation is

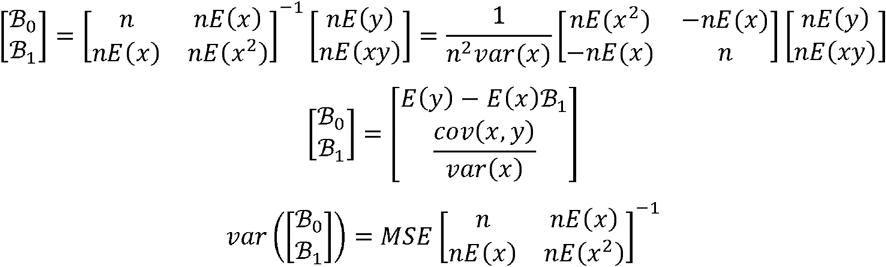

in which MSE is

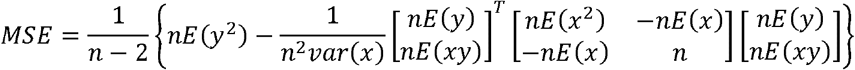

In particular 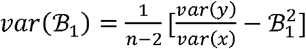

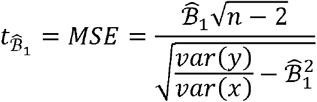

For GWAS ℬ_1 =_ *a,var*(*x)* or 1 if the locus has been standardized.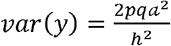.

For HE-reg ℬ_1_ is as shown in **Eq 6**. *π* takes value of 1, 0.5, and 0, with probabilities of 0.25, 0.5, and 0.25, so 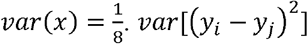 needs more calculation, please see **Appendix II**.

## Appendix II What is *var*[(*y*_1_ − *y*_2)_^2^ ]

For HE-reg, ***Y***_**12**_ ***= y***_1_ − ***y***_2)_^2^ for a pair of sibs. We unfold *var*[( ***y***_1_ − ***y***_2)_^2^] as below.

There are six unique complex terms involved, we need to treat them one by one.

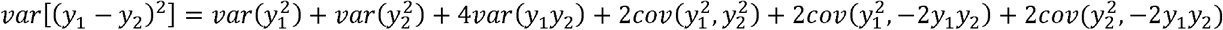

As ***y***_1_ and ***y***_2_ are related, using Cholesky decomposition, we rewrite ***y***_1_ and ***y***_2_ as below

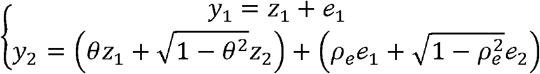

in which *θ* the relatedness score (*θ=*0.5 for a pair of sibs), *z*_1_ ∼*N*(0,*h*^2^),*e*_1_∼*N*(0,1−,*h*^2^), and *z*_2_ ∼*N*(0,*h*^2^), *e*_2_∼*N*(0,1−,*h*^2^), and *ρ*_*e*_ the correlation between residuals.

Item 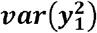:

For 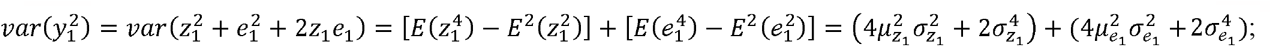 if 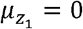 and 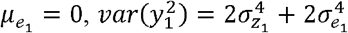

Item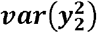:

For 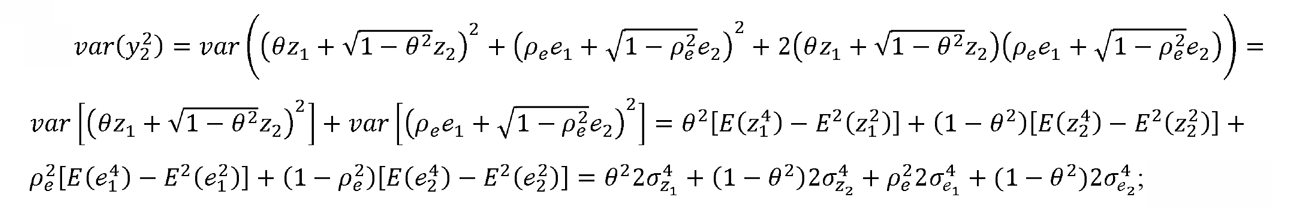 if 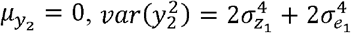.

Item ***var***(***y***_**1**_ ***y***_**2**_) :

For 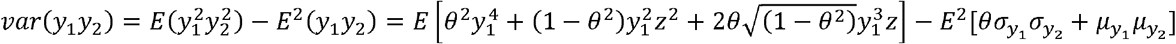

If 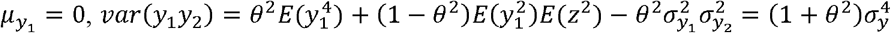

Item 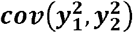:

For 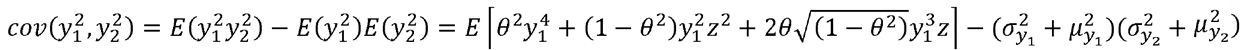

If 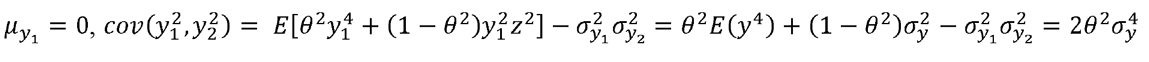

Item 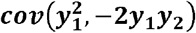:

For

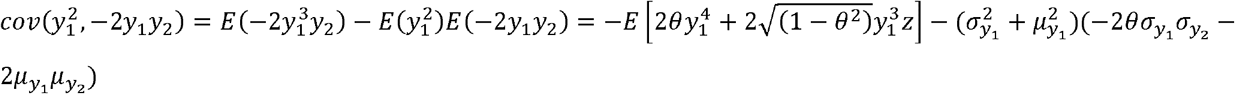

If 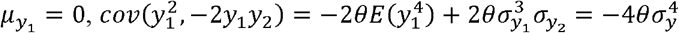

Item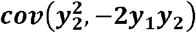:

For

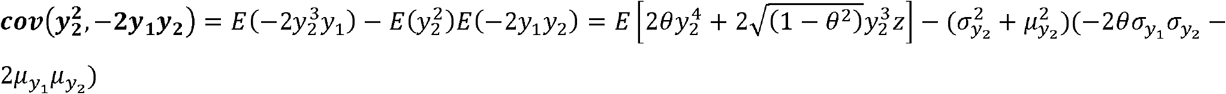

If 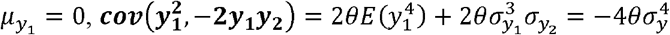

After integrating all terms together,

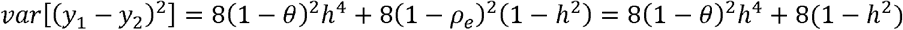

Assuming no correlation between residuals (*ρ*_*e =*_ 0, but probably it is not zero in fact).

## Appendix III IBD sharing for one allele for a full sib pair

To derive the sharing of one allele for each sib pair, we follow the general idea of Hill’s method (Hill, 1993). For a full sib pair, the IBD sharing at a locus is calculated:

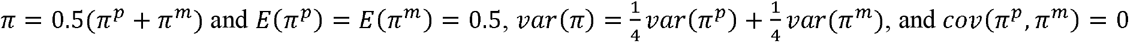 because the independency between the paternal (with superscript *p*) and maternal (with superscript *m*) meiosis. For a chromosome, with genotyped markers,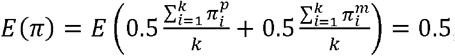, and

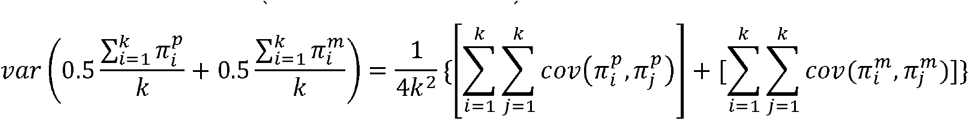

Now, we consider the IBD transmitted from paternal origins. We know, 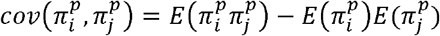, the term 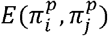 can be calculated below 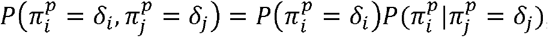, where δ =1 if the alleles are IBD for the sib pair or 0 if not.

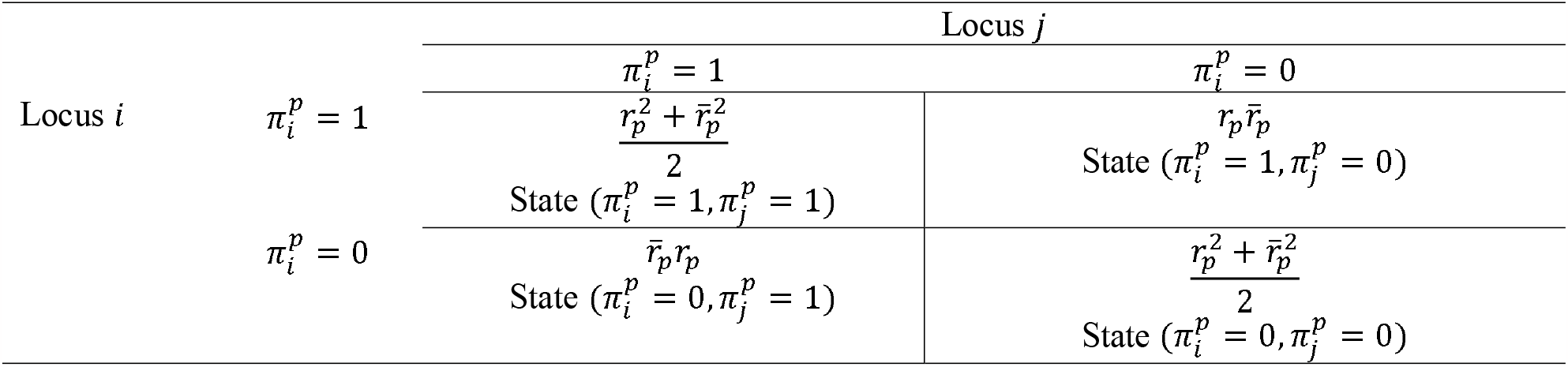

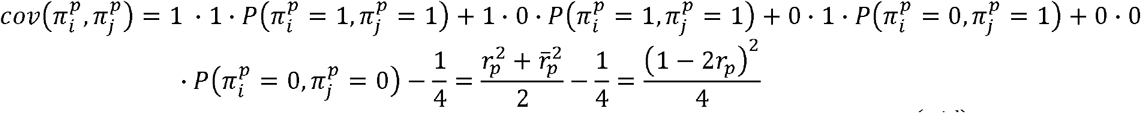

Assuming the Haldane mapping function *r*_*p*_ *=* 0.5[1−exp(−2*d*)], then, 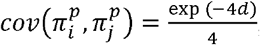 where is the genetic distance measured in Morgan. For the maternal haploids, if the recombination fractions are different from that of paternal, the similar table should be made for maternally raised IBD.

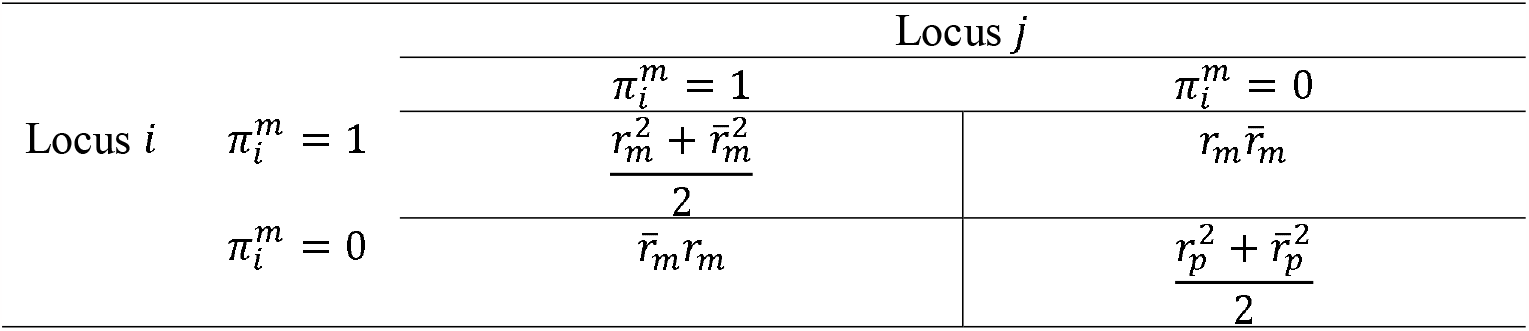

Assume the difference between maternal and paternal recombination fractions is ϵ,

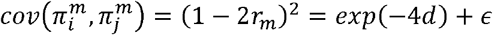

Now

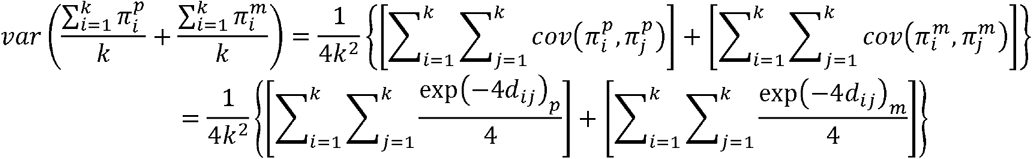

When is very large, it can be expressed as an integral, and the analytical solution is:

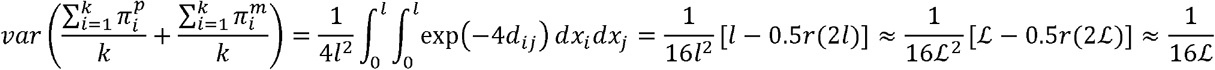

If we consider the 22 autosomes as a very long chromosome of length ℒ, and *r*(2ℒ)≈0.5 is the recombination fraction for length 2 ℒ.

